# Live Cell Lineage Tracing of Dormant Cancer Cells

**DOI:** 10.1101/2022.10.08.511405

**Authors:** Hyuna Kim, Anna Wirasaputra, Aritra Nath Kundu, Jennifer A.E. Esteves, Shelly R. Peyton

## Abstract

Breast cancer is a leading cause of global cancer-related deaths, and metastasis is the overwhelming culprit of poor patient prognosis. The most nefarious aspect of metastasis is dormancy, a prolonged period between primary tumor resection and relapse. Current therapies are insufficient at killing dormant cells; thus, they can remain quiescent in the body for decades until eventually undergoing a phenotypic switch, resulting in metastases that are more adaptable and more drug resistant. Unfortunately, dormancy has few *in vitro* models, largely because lab-derived cell lines are highly proliferative. Existing models address tumor dormancy, not cellular dormancy, because tracking individual cells is technically challenging. To combat this problem, we adapted a live cell lineage approach to find and track individual dormant cells, distinguishing them from proliferative and dying cells over multiple days. We applied this approach across a range of different *in vitro* microenvironments. Our approach revealed that the proportion of cells that exhibited long-term quiescence was regulated by both cell intrinsic and extrinsic factors, with the most dormant cells found in 3D collagen gels. We hope this approach will prove useful to biologists and bioengineers in the dormancy community to identify, quantify, and study dormant tumor cells.

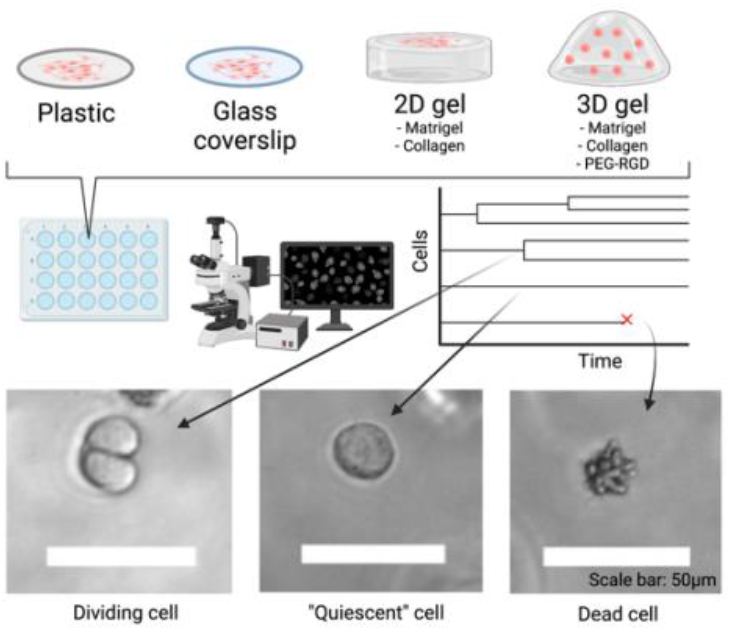

## Introduction

2.3 million women worldwide were diagnosed with breast cancer in 2020, with approximately 685,000 reported deaths.^[1,2]^ The 5-year and beyond survival rate varies based on region and breast cancer subtype, with 85-90% survival in high-income countries and 60% or lower in many developing countries.^[3,4]^ It is important to note, however, that these statistics are associated with reporting disparity from lower-middle income areas.^[5]^

When breast cancer is at stage I or II, tumor growth is controllable with chemotherapy, surgery, and radiation, and 5-year survival rate is 90-100% in the United States.^[6,7]^ However, when breast cancer metastasizes to other organs, the survival rate dramatically decreases to 22%.^[8]^ Breast cancer commonly metastasizes to bone, lung, brain, and liver. Even after initially successful treatment, 13-30% of early-stage breast cancer patients develop cancer relapse in distant organs, most frequently in the bone.^[9,10]^ This indicates that disseminated tumor cells (DTCs) can remain dormant for many years, even decades, before growing into a detectable, symptomatic tumor.^[9,10]^ It is difficult to treat these dormant cells as traditional chemotherapies target rapidly growing cells. Also, regardless of their cell cycle status, DTCs are actively protected by their microenvironment and vascular endothelium.^[11]^

Dormancy is categorized either as tumor mass dormancy or cellular dormancy. Tumor mass dormancy states that the rate of cancer cell proliferation in a bulk of tumor equals that of cell death.^[12]^ Cellular dormancy is a result of an individual DTC temporarily exiting the cell cycle and remaining in a quiescent state, with the possibility that it can resume proliferation later.^[12]^ Researchers have been studying the critical factors and the molecular mechanisms governing dormancy and subsequent relapse, but it is challenging to find and track individual DTCs that are both viable and dormant. For instance, immunohistochemistry (IHC) of fixed clinical and *in vivo* specimens can provide insight into localization of dormant or proliferating cells within the matrix by comparing the levels of Ki-67 expression.^[13,14]^ However, IHC cannot determine if the observed non-proliferative cells are capable of eventual outgrowth, nor if the factors from their microenvironment would affect the outgrowth because the process is limited to fixed samples.

Moreover, most of the traditional *in vitro* models fail to capture the interactions between cancer cells and their microenvironment. One exception is work from the Ghajar lab, who, via intravital imaging, showed that certain microenvironments, like perivascular niches, protect DTCs from chemotherapy.^[11]^ The overwhelming advantage of intravital imaging is the ability to watch DTCs in an *in vivo* context, but is expensive, difficult to learn, and is low throughput. We see intravital imaging as an excellent way to validate hypotheses, but not well-suited for hypothesis screening. Creating *in vitro* approaches to observe cell quiescence and reactivation are sorely needed to better understand the factors that lead to relapse, such as that demonstrated here.

In order to track cell plasticity with greater accuracy, Laura Heiser’s group modified an existing Markov model to better classify cells according to their heterogeneous phenotypes across lineages.^[15]^ We used this model to create cell lineage trees and analyzed individual cell proliferation and death. This method can solve the problems that other commonly used cell proliferation assays have. For example, the MTT assay is an endpoint assay, and it often overestimates cell viability. Ki-67 staining is difficult to quantify, and it requires cell fixation. Live microscopy and manual tracking of cells through this lineage tracking allowed us to find and distinguish quiescent cells from proliferating and dying cells. We present this approach here, validated with established markers of cellular dormancy, to demonstrate the emergence of dormant populations across a variety of cell culture environments.

## Results

### IHC subtypes of breast cancer cell lines correlates with observed proliferation and dormancy rates

HCC1954 is a HER2 enriched (HER2+) breast cancer cell line, which is a subtype correlated with more metastasis in the clinic, and proliferative than luminal A type cell lines such as MCF7 (from Cellosaurus). We observed the HCC1954 cell lineage trees to have a higher distribution of cells proliferating twice or more than those in MCF7 (Figs. 1a-b, S1a-b), as expected. More than 50% of the randomly selected HCC1954 cells divided twice or more (Figs. 1a-b). According to Cellosaurus, a cell ancestry database, HCC1954 has a doubling time of 45 hours and MCF7 has a maximum doubling time of 80 hours, which further validates the difference in cell proliferation between the two cell lines. When comparing the cell lineage trees of HCC1954 and MCF7 cultured on tissue culture polystyrene (TCPS), there are a smaller number of non-dividing cells, illustrated as a single straight line across the graph, in HCC1954 than in MCF 7 (Figs. 1a-b, S1a-b). In this paper, we define the non-dividing live cells as “dormant” cells, as they don’t divide and don’t die but rather persist over the entire course of the experiment.

**Fig. 1.**
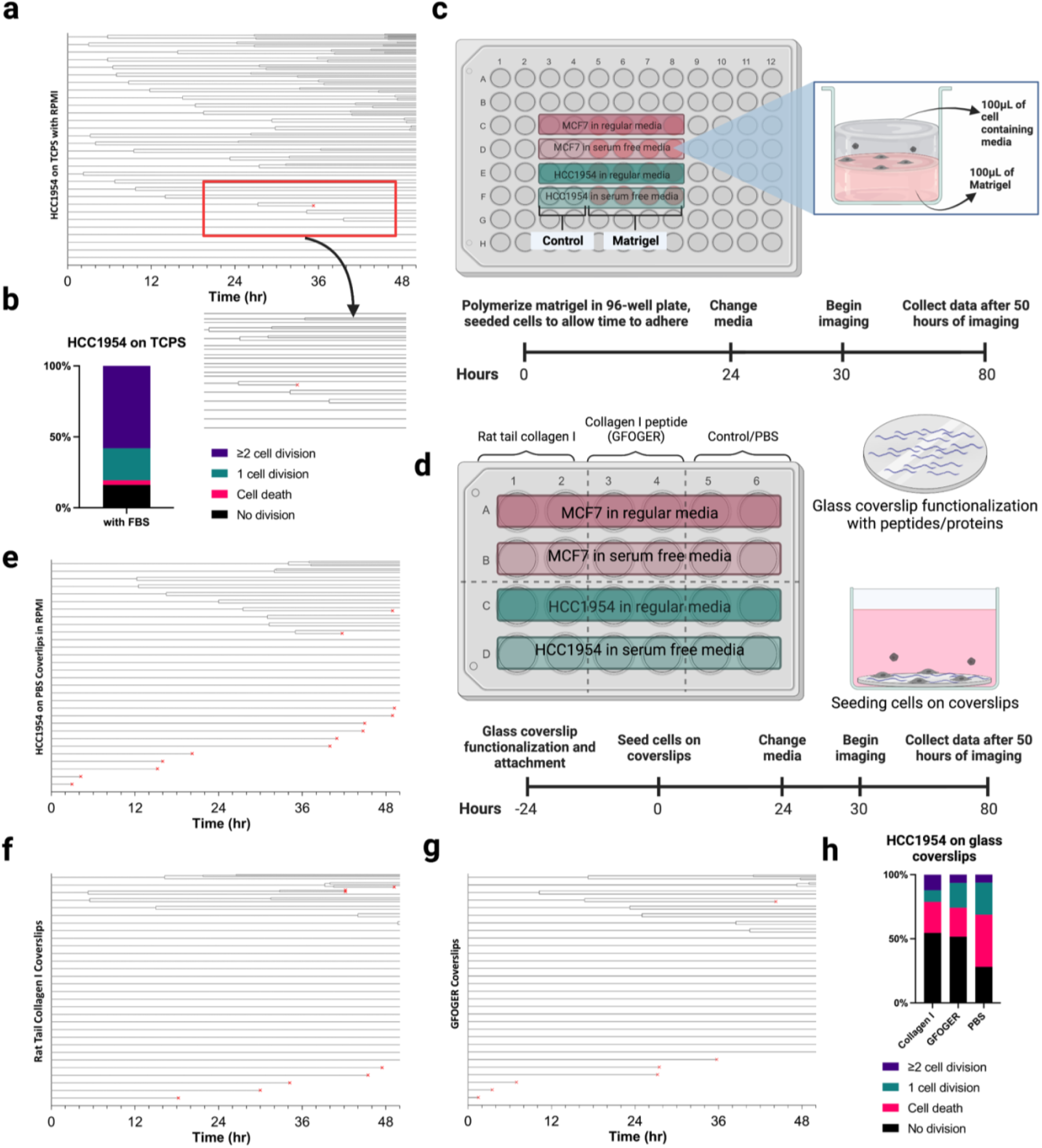
Experimental setups and timelines for 2D environments. (**a**). Shows a cell lineage tree of HCC1954 cells on TCPS obtained by randomly tracking 30 individual cell proliferation across 50 hours. Each time a cell splits, it is denoted in the graph by splitting into two lines. Increasing on the y-axis shows least proliferating to most. If a cell doesn’t split, it will have a straight line across the entire graph. When encountering a dying cell, the time of death is marked with a red ‘x’. (**b**). Bar graph distribution of cell division and death from the lineage analysis. (**c**). This schematic consists of the seeding procedure and experimental timeline of two cell lines (MCF7 and HCC1954) in different media conditions on top of 2D Matrigel in a 96-well plate. (**d**). Schematic of a 24-well plate containing protein treated coverslip conditions with an included timeline. (**e-g**). Cell lineage analyses of HCC1954 in serum containing media (RPMI) across the coverslip conditions: (**e**). PBS-treated coverslip, (**f**). Collagen I protein treated coverslip, and (**g**). Collagen I peptide (GFOGER) treated coverslip. (**h**). Stacked bar graph for coverslip conditions from figures e-g.

### Dormant cell population increases on glass coverslips compared to that on TCPS

We tracked cells on glass substrates that were chemically functionalized with individual cell proteins or peptides, as we have described elsewhere.^[16,17]^ Cell attachment was overall lower on these protein- and peptide-functionalized glass coverslips, and we also observed higher rates of cell death than those on TCPS (Figs. 1a-b, 1e-h). However, the number of dormant cells was significantly higher on glass coverslips than on TCPS. Overall, the dormant cell population on both the rat tail collagen I functionalized and the collagen I peptide (GFOGER) functionalized coverslips had similar trends (Figs. 1f-h); though, the cells seeded on the GFOGER functionalized coverslips had a lower number of cells that divided twice or more than the cells seeded on the collagen I protein functionalized coverslips (Fig. 1h). Additionally, cells cultured on the protein- or peptide-functionalized coverslips had significantly higher portion of dormant cells while had lower cell death than the control, PBS-treated coverslips (Figs. 1e-h). Together, this tells us that collagen-functionalized glass coverslips allow cells to undergo dormant, and the form of collagen – either whole collagen protein or a part of the protein which is GFOGER peptide – does not have a significant effect on dormant cell population.

### The effect of naturally derived matrix materials on dormant cell population is significantly higher in 3D than on 2D

Matrigel and collagen gels are commonly used materials for cancer studies as collagen is the most abundant protein in human connective tissues, and Matrigel is extracellular matrix (ECM) extracted from Engelbreth-Holm-Swarm mouse tumors.^[18]^ When MCF7 and HCC1954 cell lines were cultured on 2D Matrigel, both were less proliferative in comparison to TCPS (Figs. 1b, 2b, S1, S2). The HCC1954 line had a greater number of cells trackable (at time 0); however, had a greater number of deaths over the course of the experiment for both serum-containing and serum-free media conditions (Fig. 2b). This suggests those cells have a greater sensitivity to serum deprivation, which we also observed in a previous dormancy study.^[19]^ Additionally, both cell lines cultured on 2D Matrigel appeared to have lower proliferation than when cultured on TCPS, which could be due to the higher softness of Matrigel, which is known to influence cell proliferation rates.^[20,21]^ However, the dormant cell population does not significantly change in both cell lines on 2D Matrigel compared to those on TCPS (Figs. 1a-b, 2b-c, S1, S2a-b). MCF7 cells showed less cell death on 2D Matrigel (Figs. 2c, S2a) while HCC1954 suffered more cell death (Figs. 2b, S2b). Also, there were more dormant MCF7 cells cultured for 50 hours (Figs. 2c, S2a) than HCC1954 (Figs. 2b, S2b), in both serum-containing and serum-free growth media conditions. Together, 2D Matrigel environment does not have a significant effect on dormant cell populations in both HCC1954 and MCF7, but the dormant cell population differs by cell line, which means the IHC subtypes of breast cancer cell lines play more significant role in dormancy than the 2D Matrigel environment itself does.

**Fig. 2.**
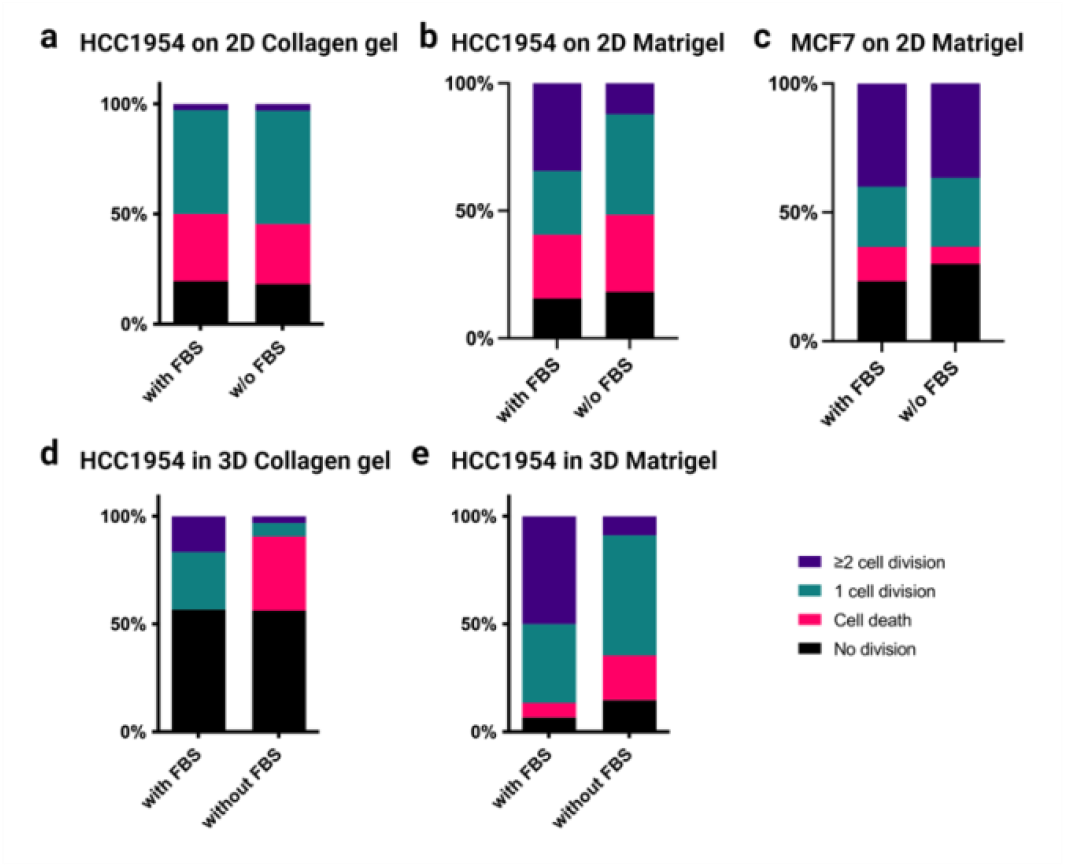
Distribution of cell proliferation in 2D and 3D gel environments. (**a**) The stacked bar graph showing proportion of cells with 0, 1, 2+ cell divisions or with cell death, which data was taken from the cell lineage analysis conducted on HCC1954 cells seeded on 2D collagen gel in regular RPMI. (**b-c**). Bar graph showing proportion of cells with 0, 1, 2+ cell divisions or with cell death for (**b**) HCC1954 and (**c**) MCF7 cells on 2D Matrigel. (**d**). Bar graph showing proportion of cells with 0, 1, 2+ cell divisions or with cell death for HCC1954 cells in 3D collagen gel. (**e**). Bar graph showing proportion of cells with 0, 1, 2+ cell divisions or with cell death for HCC1954 in 3D Matrigel.

Significant differences in cell growth behavior, especially the dormant cell population, were observed when compared these cell lines on the same ECM protein in 2D versus 3D. HCC1954 cells were cultured on 2D collagen gels and in 3D collagen gels, and more than 50% cells were dormant in 3D collagen gel environment (Figs. 2d, S3b) while less than a quarter of the cells were dormant when cultured on a 2D collagen gel (Figs. 2a, S2c). Interestingly, about the same number of cells were dormant for 50 hours in both serum-containing and serum-free conditions in 3D collagen gels, but there were greater numbers of cells dying in the 3D collagen material than 2D (Figs. 2d, S3b). While the 3D collagen environment resulted in a higher number of dormant cells than 2D collagen gel environment, it showed the opposite trend in Matrigel. There were a greater number of dormant cells on 2D Matrigel (Figs. 2b, S2b) than in 3D Matrigel (Figs. 2e, S3a). Additionally, when comparing the matrix materials in the same dimension, 2D collagen gel vs. 2D Matrigel for example, the matrix material does not play a significant role in dormancy when it’s on 2D (Figs. 2a-b); however, when it’s in 3D, there was a significant difference in dormant cell population between cells in 3D collagen gels and those in 3D Matrigel. These results show that the types of matrix materials have a greater effect in 3D than on 2D.

### Immunofluorescence staining validates the cell lineage tree approach to identify dormant cells

Two different mouse breast cancer cell lines from the same origin were used to validate if the cell lineage tree analysis is a viable method to use for dormancy study. D2A1 is a proliferative cell line and D2.0R is a dormant cell line.^[22–25]^ Both cell lines showed high proliferation in serum containing medium in 3D Matrigel (Figs. 3a-b). Significant differences in proliferation and dormancy between these two cell lines was observed when they were cultured in serum-free media. D2.0R, the dormant cells, showed less cell death and more dormant phenotypes in serum-free environment than D2A1, the proliferative cells, and there were more non-dividing alive D2.0R cells when cultured in serum-free media than in serum-containing media (Figs. 3a-b). This result supports an earlier study that showed that serum deprivation forced cells into dormant-like state.^[19]^

**Fig. 3.**
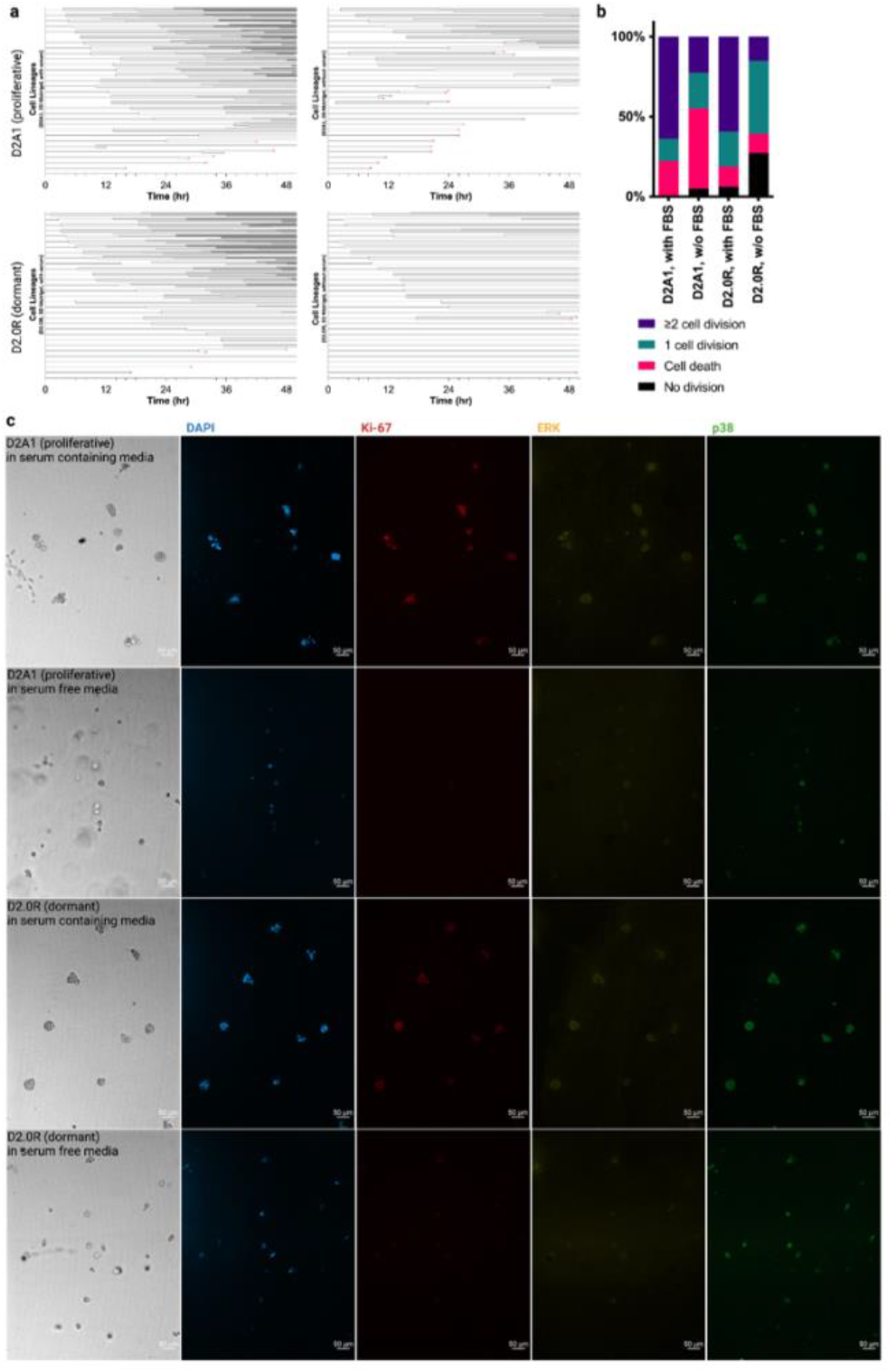
Mouse breast cancer cells that have proliferative (D2A1) or dormant (D2.0R) phenotype in 3D Matrigel. (**a**). Cell lineage trees of D2A1 (proliferative) and D2.0R (dormant) cells in serum containing and serum free conditions. Top row is showing D2A1 lineages, and bottom row is showing D2.0R lineages. Left column is the lineage trees of cells under serum containing condition, and right column is the lineage trees of cells under serum free condition. (**b**). A stacked bar graph showing proportion of cells with 0, 1, 2+ cell divisions or with cell death of D2 series cells, and the number of each group was calculated from the cell lineage trees shown in (**a**). (**c**). IF staining images of D2 series cells in serum containing and serum free conditions. Images on the far-left column are brightfield images, and the images from 2nd to 5th columns are IF staining images. Cells were stained with DAPI (blue), Ki-67 (red), ERK (yellow), and p38 (green). Scale bar = 50μm.

Cells were immunofluorescently stained with Ki-67, ERK, p38 and DAPI to validate that the differences we saw in numbers of dormant cells using cell lineage tracking was reflected by the standard set of markers used to identify dormant cells. Dormant cells are distinguished by low Ki-67, low ERK, and high p38 expression.^[26,27]^ No significant difference was observed in the stained markers, and there were low number of dormant cells in serum containing environment for both cell lines, D2A1 and D2.0R (Fig. 3c, first and third rows). There were a smaller number of cells due to high cell death in D2A1 cell culture in serum free condition (Fig. 3c, second row) than D2.0R cells (Fig. 3c, fourth row). The D2.0R cell culture in serum free condition in 3D Matrigel had the most number cells stained for the dormancy markers than any other condition (Fig. 3c), validating our results from the cell lineage tree analyses.

### Applications in 3D synthetic hydrogels

Several synthetic *in vitro* platforms have been developed to compete with naturally derived materials like collagen gel and Matrigel, as the synthetic materials enable more tunability and more precise control over its properties than protein-based hydrogels.^[18]^ We cultured HCC1143, triple negative breast cancer cells, in 3D poly(ethylene glycol) (PEG) hydrogels containing widely used integrin-binding peptide sequence Arg-Gly-Asp (RGD) and partially crosslinked with a matrix metalloproteinase (MMP) sensitive peptide. Cells proliferated more in the degradable PEG-RGD hydrogels than in the non-degradable hydrogels. Less cell death and faster cell growth were achieved in growth media condition supplemented with epithelial growth factor (EGF) than in regular serum containing media, but the dormant cell population does not change significantly upon EGF addition (Figs. 4a, S4a). This suggests that the addition of growth factors does influence cell proliferation, but it does not play a significant role in dormant cell populations in 3D synthetic PEG-RGD hydrogels. Rather, the presence of MMP-degradable sites in the 3D synthetic environment plays more significant role in dormant cell populations.

**Fig. 4.**
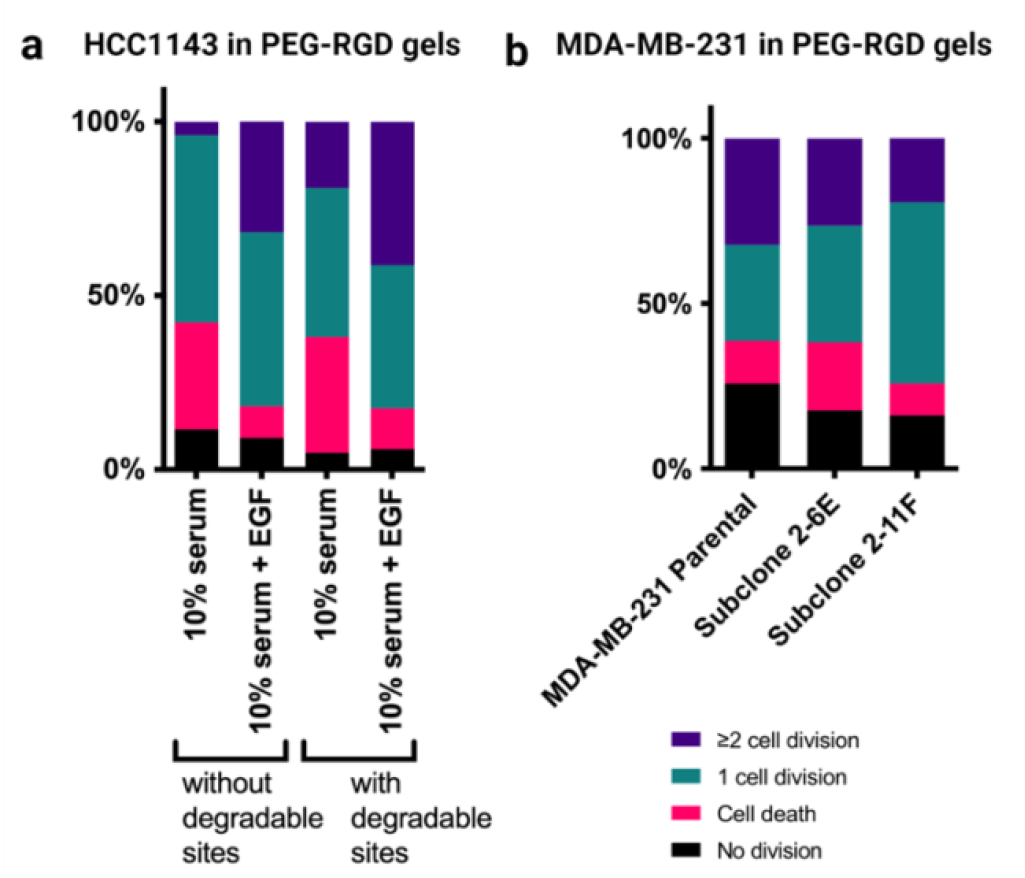
Triple negative breast cancer lines in PEG-RGD gels. (**a**) A stacked bar graph from the cell lineage analysis conducted on HCC1143 cells in a 3D PEG-RGD system with different serum and EGF conditions (**b**). Bar graph of the MDA-MB-231 cell line with its subclones 2-6E and 2-11F in 3D PEG-RGD gels.

### Cell clonality differences in dormancy tracking

Thus far, all studies were performed on heterogeneous cell lines, where observations could be determined by both microenvironment and the genetic differences within these populations. Thus, we created cell subclones from the otherwise heterogeneous MDA-MB-231 cells, a triple negative breast cancer cell line,^[28]^ and its cell subclones were cultured in PEG-RGD hydrogels of 2 kPa modulus with MMP-degradable sites. Although the subclones originated from the same cell line, their proliferation, death, and dormancy rates varied significantly, demonstrating the power of cell intrinsic factors in driving dormancy and growth (Fig. 4b). MDA-MB-231 parental cells showed the highest number of dormant cells, which also reflects our previously published observation.^[19]^ Together, these results conclude that the cell lineage tree analysis can be used to analyze growth behavior of cells of various subtypes on diverse culturing environment to track and identify individual dormant cells while keeping them alive.

## Discussion

Currently, there is no set method to monitor dormant states or predict the probability of recurrence from these cell populations—there are many biological models that provide some depth in understanding these characteristics of dormancy behavior; however, many are still pre-clinical and lack accuracy.^[29]^ In addition, the bulk of the field’s knowledge of tumor and cellular dormancy is limited to static, endpoint measurements *in vivo*. Using the cell lineage tree analysis, we demonstrated that we can track individual cells while keeping them alive, whereas other types of immune-staining is an endpoint assay where dormant cells can be distinguishable by marker expressions.

The biggest difference we observed between the two breast cancer cell lines used (MCF7 and HCC1954) is the individual sensitivity when subjected to serum-free media conditions. Some studies have shown that it is possible to grow certain breast cancer lines without serum in their growth media.^[30]^ This is important because successful serum-free growth conditions will aid in better biological studies without the interference of hormones or other serum proteins which differ by batch. Our results show that there is little change in terms of cell death when comparing the growth conditions between the 2D environments.

It is known that biological subtypes of breast cancer are associated with the rate and the region of recurrence.^[31,32]^ A study^[31]^ showed that TNBC had the highest rate of recurrence in the first five years, whereas the recurrence rate for luminal A and luminal B tumors was initially low but remained continuously even after 10 years of follow-up. Another study showed that among several breast cancer subtypes, estrogen receptor positive (ER+) breast cancer is most commonly associated with bone metastasis and dormancy, relative to the other breast cancer subtypes.^[33]^ For the different subtypes, recurrent breast cancer is normally associated with the luminal A group, which can usually be identified by the estrogen positive. This is the most prevalent subtype of cancer, and typically results in better prognosis. We observed that the MCF7 cells were more sensitive to serum conditions than the HCC1954 cells, as there is no significant difference in dormant cell populations for the latter (Figs. 2b-c); thus, we speculate that the ER+ cell line has a greater sensitivity to serum conditions than the HER2+ group in terms of the dormant cell population.

After successful extravasation, DTCs can undergo decades of dormancy in secondary metastatic sites— bone marrow, liver, lung, and brain. There are two types of dormancy: cellular dormancy and tumor dormancy. Cellular dormancy is when individual cells are not proliferative enough to be detectable, and tumor dormancy is when there is an equilibrium between cell proliferation and death across the bulk of the tumor mass.^[34]^ DTCs can enter a largely quiescent state with no proliferative growth (cellular dormancy) before reactivating under a phenotypic switch. These reactivated cells are often more invasive and drug-resistant relative to their parental cells, resulting in diminishing patient prognosis. In our study, we focused on cellular dormancy by seeding at a low enough density to more accurately trace the lineage of single cell populations to eventually study what microenvironmental factors are most likely to generate dormant cells.

In our study, we have cell lines from the following subtypes: luminal A, HER2+, and triple negative with increasing aggression, respectively. We also tested mouse breast cancer cell lines, D2A1 (proliferative) and D2.0R (dormant). We found that breast cancer cells with different subtypes show distinct proliferating behavior; some died less and proliferated better in serum-free conditions than other subtypes, and some showed more quiescent phenotype than others in serum-free environment. Also, mouse breast cancer D2 cell lines were overall proliferating much faster than human breast cancer cells.

Together, our results show that there are multiple factors that affect cell growth and dormancy. Breast cancer cell types are one of the factors, but the cell culture material and dimensionality also have a great effect on the cell proliferation and dormancy. It aligns with a previous study by Kloxin group; Different cell subtypes showed distinct dormancy scores in a same microenvironment that they developed for long term 3D dormancy culture model.^[35]^ In their synthetic matrix, ER+ breast cancer cells underwent dormancy while triple negative breast cancer cells did not.^[35]^ The conclusion that the culturing matrix and dimensionality also play roles in cell growth behavior, is supported by an earlier study that describes cancer cells showed distinct phenotype and marker expressions depending on culturing environment.^[36]^ Specifically for dormant cell behaviors, there are multiple studies that showed the degradability of materials or immobilization by environment could induce cellular dormancy.^[37,38]^ For example, hydrogels with no adhesivity but high degradability induced cellular dormancy, while hydrogels with high adhesivity and degradability promoted growth.^[37]^ Also, non-degradable hydrogels immobilize cells in the microenvironment and dormancy-capable cells can become dormant in such system due to the physical confinement.^[38]^

In sum, the cell lineage tree approach is a good model to study single cell heterogeneity^[15]^ and this analysis method can be applied to variety of cells with various culturing environment. Gross *et al*. used the cell lineage tree model to capture cell cycle dynamics upon drug application, which includes drug-specific effects on cell cycle phase and single cell responses.^[39]^ One of the advantages of this method is that we can trace live cells real time unlike other endpoint assays. This will help us to better understand cellular dormancy and how it is related to the tumor microenvironment of DTCs.

## Materials and Methods

### Breast cancer cell culture

Different types of human breast cancer cell lines, including MCF7, MDA-MB-231, HCC1954, and HCC1143 were used for this study. MCF7 and MDA-MB-231 cells were cultured under 5% CO2 and 37ºC with DMEM containing 10% fetal bovine serum (FBS) and 1% penicillin/streptomycin (pen/strep). MDA-MB-231 cell subclones were generated by Ning-Hsuan Tseng,^[28]^ and the same culturing method was used as the parental MDA-MB-231 cell line. HCC1954 and HCC1143 were cultured under 5% CO2 and 37ºC with RPMI containing 10% FBS and 1% pen/strep. Two different types of mouse mammary tumor cells were also used, which were derived from spontaneous mouse mammary tumors, originated from D2 hyperplastic alveolar nodule in Miller lab.^[40,41]^ D2A1 (proliferative) and D2.0R (dormant) cells were cultured under 5% CO2 and 37ºC with DMEM containing 10% FBS and 1% pen/strep.

### Live cell microscopy for tracing individual cells

Cells were seeded on TCPS, functionalized glass coverslips, 2D collagen gel, 2D Matrigel, or in 3D collagen gel or 3D Matrigel. The cell culture media was changed 24 hours after cell seeding, to either normal growth medium (with 10% FBS) or serum free medium. Six hours after the media change, time-lapse images of single cells were taken for 50 hours, at either 15-minute (for 2D experiments) or 30-minute intervals (for 3D experiments). Imaging was performed using Zeiss Axio Observer Z1 Inverted Fluorescence Microscope.

### Preparation of culturing environment and cell seeding for 2D environment

For tissue culture polystyrene (TCPS): After at least 80% confluence achieved in T-25 flasks, we harvested cells for seeding on different substrates with varying media conditions. Used approximately 2mL of 1X phosphate buffered saline (PBS) to wash the flask before using 2mL trypsin for 2-5 minutes to cleave cells off flask surface. Then we deactivated trypsin with 4mL of regular media. Placed the gathered liquid and cells in a 10mL conical tube in a centrifuge for 5 minutes at 200G. Aspirated the cell pellet by vacuuming the supernatant and resuspend the cells in 1mL of media. Counted the cells using an automated cell counter (a hemocytometer has been used with near similar results) and mathematically calculated the volume needed to seed approximately 5000 cells in a 6-well plate. Used similar ratios for greater-numbered plates by dividing the cells with the surface area given by Thermo Fischer and other companies. After seeding, we waited 24 hours for cells to adhere before changing media conditions (serum free DMEM and RPMI media). After the media change, we waited an additional 6 hours before beginning imaging for 50 hours.

For 2D Matrigel: We thawed Matrigel (Corning) overnight in the 4_-_°C fridge to prepare for use the following day. 80% confluence was achieved in T-25 flasks, harvested the cells to use with different growing medium on top of the Matrigel. Once completely sterilized, 100µL Matrigel was slowly pipetted into a warm 96-well plate; then was placed to polymerize in the 37°C incubator for 20 minutes. While waiting for the Matrigel to solidify, the same cell harvesting procedure as described in TCPS method was used. Calculated a 200-cell density in each well placed in 100µL of media to seed on top of the Matrigel layer. Waited 24 hours for cells to adhere before changing media conditions and imaging (Fig. 1c).

For 2D collagen gel: Rat tail collagen I was taken out of 4ºC fridge and kept on ice. 410μL of rat tail collagen I solution at 3mg/mL was mixed with 520μL of serum free media and 25μL of sterile filtered 1M NaOH in microcentrifuge tubes, final concentration of which is 1.3mg/mL. 100μL of this mixture solution was taken and placed into a pre-warmed 96-well plate, then was placed to polymerize in the 37ºC insulator for 30 minutes. While waiting for the collagen gels to polymerize, the same cell harvesting procedure as described in TCPS method was used. 200 cells per well were seeded on top of each 2D collagen gel in a 96-well plate, and 100μL of media was placed on top. We waited 24 hours before changing media with or without serum and imaging.

For 2D functionalized coverslips: Rat tail collagen I protein and collagen I integrin-binding peptide were immobilized on glass coverslips. Briefly, 15mm coverslips were oxygen plasma treated (Harrick Plasma, Ithaca, NY, USA), and silanized through vapor phase deposition of (3-aminopropyl)triethoxysilane (Sigma-Aldrich) at 90°C for a minimum of 18 hours. The coverslips were sequentially rinsed in toluene (Fischer Scientific), 95% ethanol (Decon Laboratories, King of Prussia, PA, USA), and water, and dried at 90°C for one hour. These were subsequently functionalized with 10 g/L N,N,-disuccinimidyl carbonate (Sigma-Aldrich) and 5% v/v diisopropylethylamine (Sigma-Aldrich) in acetone (Fischer Scientific) for two hours. Finally, coverslips were rinsed three times in acetone and air-dried for 10 minutes. Rat tail collagen I protein and collagen I integrin-binding peptide were then covalently bound to the glass coverslips via the reactive amines at 2 µg/cm^2^ concentration. The peptide sequence CGPGPPGPPGPPGPPGPPGFOGERGPPGPPGPPGPPGPP (GFOGER) was used as the collagen I binding peptide. The rat tail collagen I protein and GFOGER peptide peptide were subsequently blocked using methyl-PEG24-amine (MA-PEG24) (Thermo Fisher Scientific). Post functionalization, the coverslips were UV sterilized for minimum of 1 hour. The cells were then added on the coverslips at a cell density of 2000-3000 cells/cm^2^. The cells were allowed to adhere on the surface before changing the media and imaging (Fig. 1d).

### Preparation of culturing environment and cell seeding for 3D environment

For 3D Matrigel: We thawed Matrigel overnight in the 4ºC fridge, and placed the Matrigel on ice on the day of experiment. For one gel, 5000 cells were mixed with 10μL of Matrigel, and the 10μL of Matrigel was placed on a pre-warmed 24-well plate, and the plate was put in the 37ºC incubator for 20 minutes to fully polymerize Matrigel. Once it gets polymerized, 1mL of media was put in each well.

For 3D collagen gel: Rat tail collagen I was taken out of 4ºC fridge and kept on ice. 41μL of rat tail collagen I solution at 3mg/mL was mixed with 52μL of serum free media and 2.5μL of sterile filtered 1M NaOH in microcentrifuge tubes, final concentration of which is 1.3mg/mL. 90μL of this mixture solution was taken out to resuspend cell pellets of 45000 cells for 9 collagen gels. 10μL of this solution was used to make one collagen gel with 5000 cells per gel in a 24-well plate. Once it gets polymerized, 1mL of media was put in each well.

For 3D PEG-RGD gel: 20kDa 4-arm PEG-maleimide (JenKem Technology) was dissolved in serum free media at pH7.4 at 20 wt%, mixed with 2mM RGD peptide (peptide sequence: GRGDSPCG) purchased from GenScript, and we let them react at room temperature for 10 minutes. RGD peptides were used as integrin binding peptides. 1.5kDa PEG-dithiol (JenKem Technology) was dissolved in 1X PBS at pH 7.4. The weight of PEG-dithiol was calculated to have 1:1 molar ratio of thiol to maleimide when mixed together. PEG-dithiol solution was mixed with 25 mol% of MMP-degradable peptide, GCRDGPQGIWGQDRCG, purchased from GenScript for the PEG-RGD gels with degradable sites. 1μL of PEG-dithiol solution was placed first in the middle of the well on a 24-well plate, followed by an addition of 9μL of PEG-maleimide solution with peptides and cells on top of the 1μL PEG-dithiol solution. 5000 cells per gel was seeded in each 10μL gel, and 1mL of media was put in the 24-well plate after letting the gels to fully polymerize for 5-10 minutes in the 37ºC incubator.

### Lineage Tree Analysis

After the 50-hour imaging was done in the Zeiss Axio Observer Z1 Inverted Fluorescence Microscope, all the microscope image files were transferred to open in Imaris software. Microscope files were then converted to movie files showing a timeline (Supplemental video S5). We manually marked specific time points when a cell divides or dies on an Excel spreadsheet for randomly selected cells for each condition, and generated cell lineage trees to illustrate individual cell growth.

As seen in Fig. 1a, each horizontal line indicates a single cell during the multi-hour time-lapse microscopy imaging. When a cell divides, the horizontal line diverges at that time point. When a cell dies, it is marked with a red “x” at the end. Single horizontal lines that do not diverge throughout the whole time of imaging, mean that these cells did not divide but are still alive for the entire period of time-lapse microscopy imaging.

From each cell lineage tree, we counted the numbers of non-dividing and dying cells, as well as the number of cells that divide once or that divide two or more times during the 50-hour imaging. We created stacked bar graphs in Prism software, visualizing the proportion of cells in each category (no division; cell death; 1 cell division; ≥2 cell division) (Fig. 1c).

### Immunofluorescence staining and imaging

After the 50-hour imaging is finished, cells were fixed with 10% formalin. Cells were then permeablized and stained with DAPI, Ki-67 (ab156956, Abcam), ERK (ab54230, Abcam), and p38 (#4511, CellSignaling). IF stained cells were imaged using Zeiss Cell Observer SD Spinning Disc Confocal Microscope.

## Acknowledgements

The authors would like to thank Laura Heiser and Sean Gross for helping with learning the cell lineage tracing approach. The authors would also like to thank Ning-Hsuan Tseng for providing the cell subclones of MDA-MB-231 cell line. This work was supported by a grant from the Jayne Koskinas Ted Giovanis Foundation for Health and Policy, an NSF CAREER grant (DMR-1454806), and grants from the NIH (R21 CA223783, DP2 CA186573, and U01CA265709) to S.R.P. S.R.P. was supported by an Armstrong Professorship at UMass Amherst.

## Supplementary Information and Figures

**S1.**
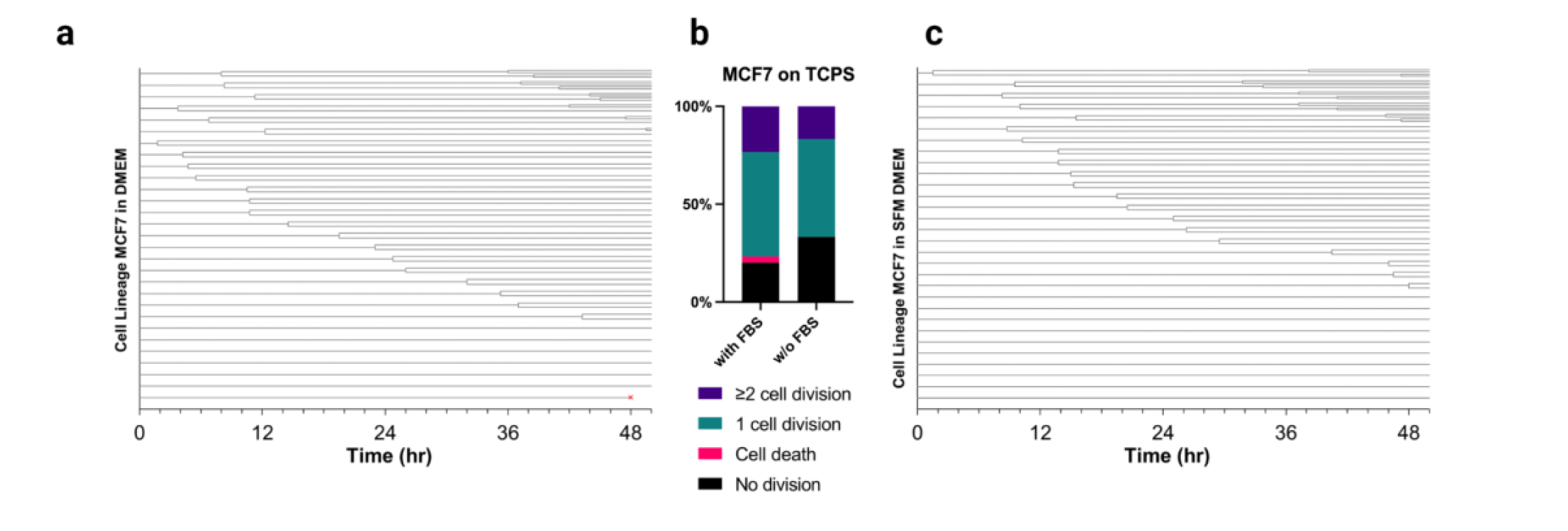
Cell proliferation data for MCF7 cell line on TCPS. Shows the proliferative capacity of MCF7 by tracking the randomly selected 30 individual cells in (**a**) serum containing and (**c**) serum free media. (**b**). Shows the proportion of cells with 0, 1, 2+ cell divisions or with cell death in a stacked bar graph.

**S2.**
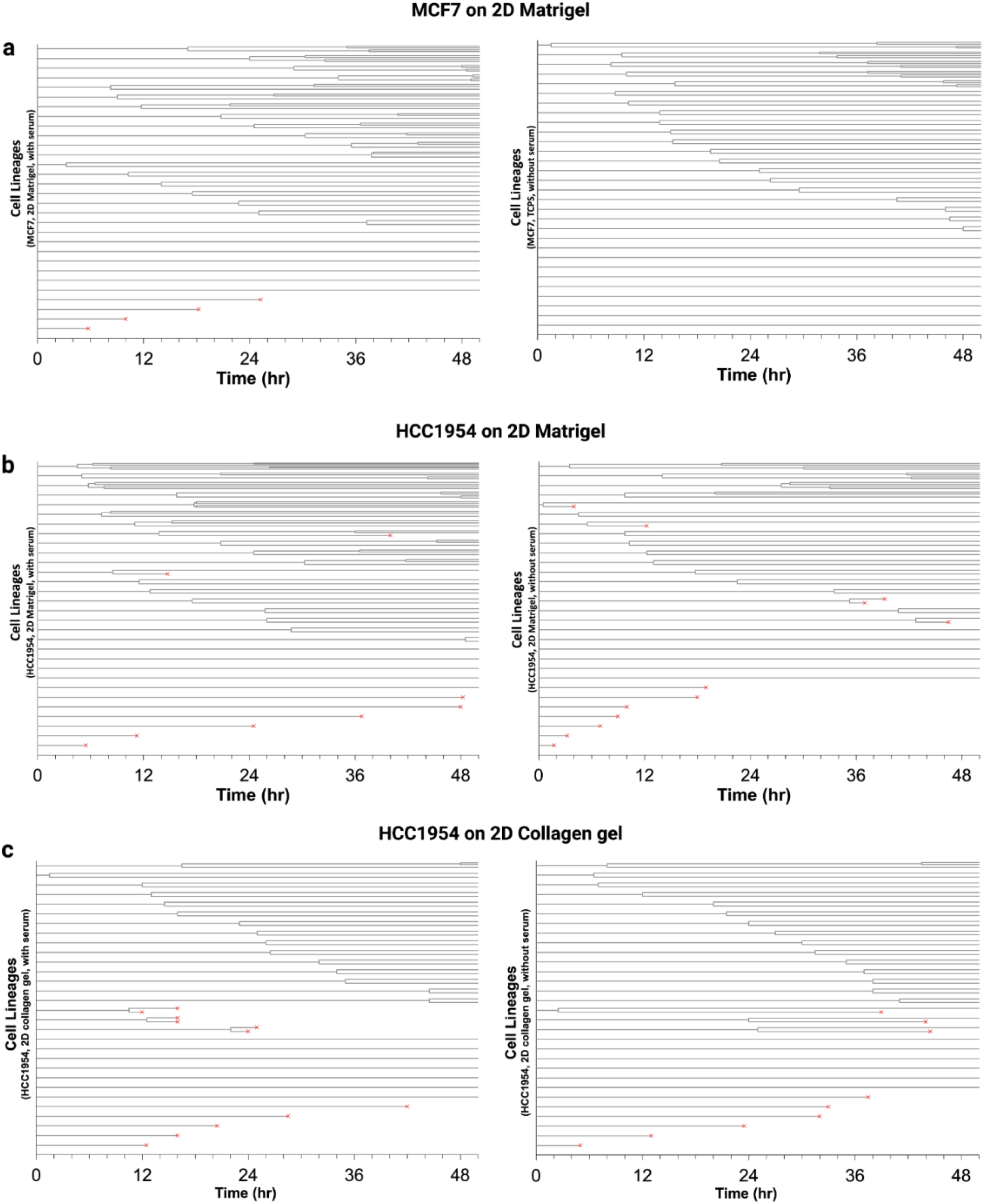
Cell lineage analysis for MCF7 and HCC1954 on 2D gel systems. (**a**). Shows the MCF7 cells cultured on 2D Matrigel for 50 hours in serum containing and serum free conditions. (**b**). Shows HCC1954 cells on Matrigel with the same conditions as (**a**). (**c**) Shows HCC1954 cells on 2D collagen gel with the same serum conditions.

**S3.**
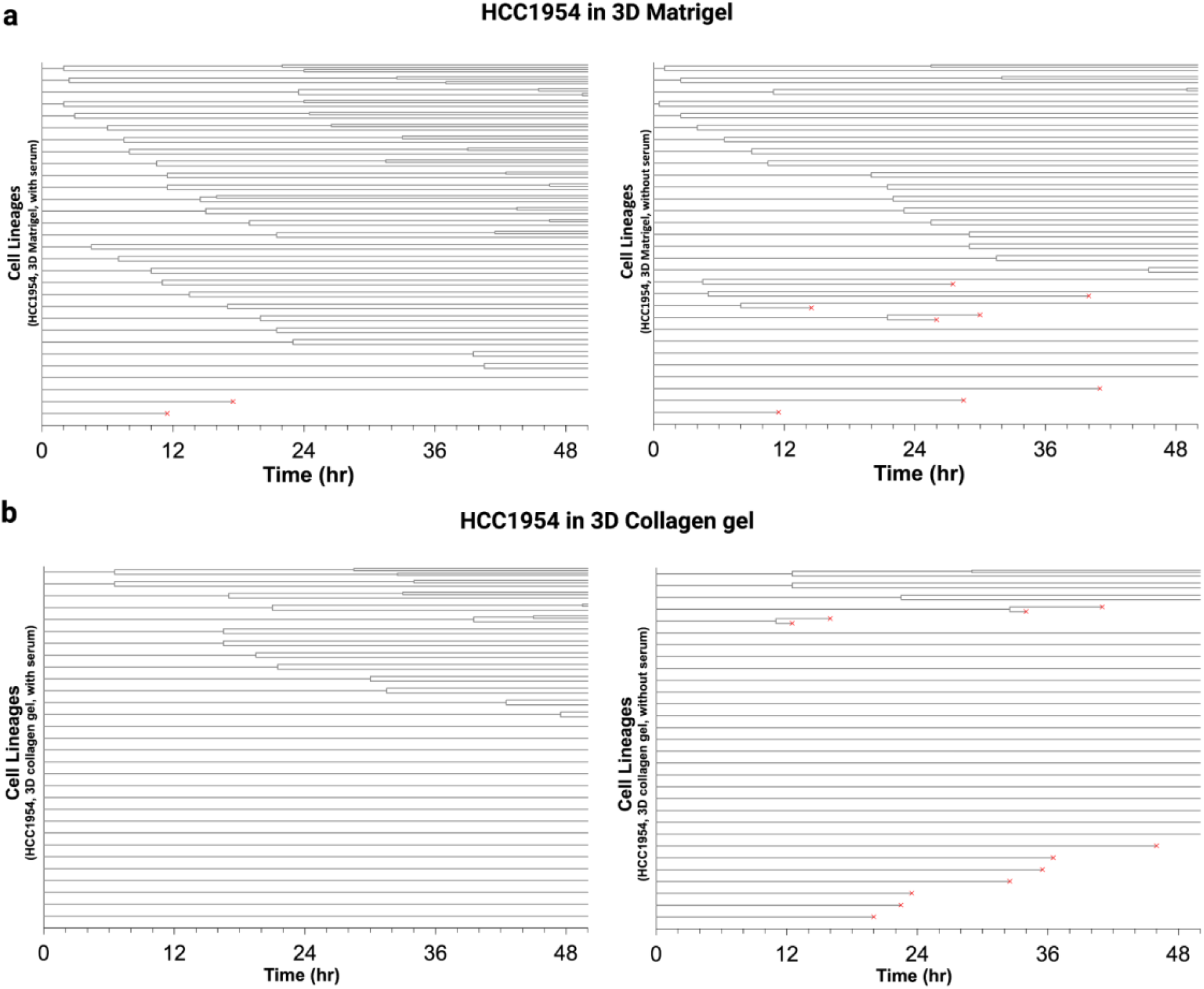
Cell lineage analysis for HCC1954 cells in a 3D gel environment. (**a**). Shows lineage analysis for HCC1954 cells cultured in 3D Matrigel for 50 hours in serum containing and serum free conditions. (**b**). Shows the HCC1954 cells cultured in a 3D collagen gel environment. 30 cells were randomly selected for (**a**) and (**b**).

**S4.**
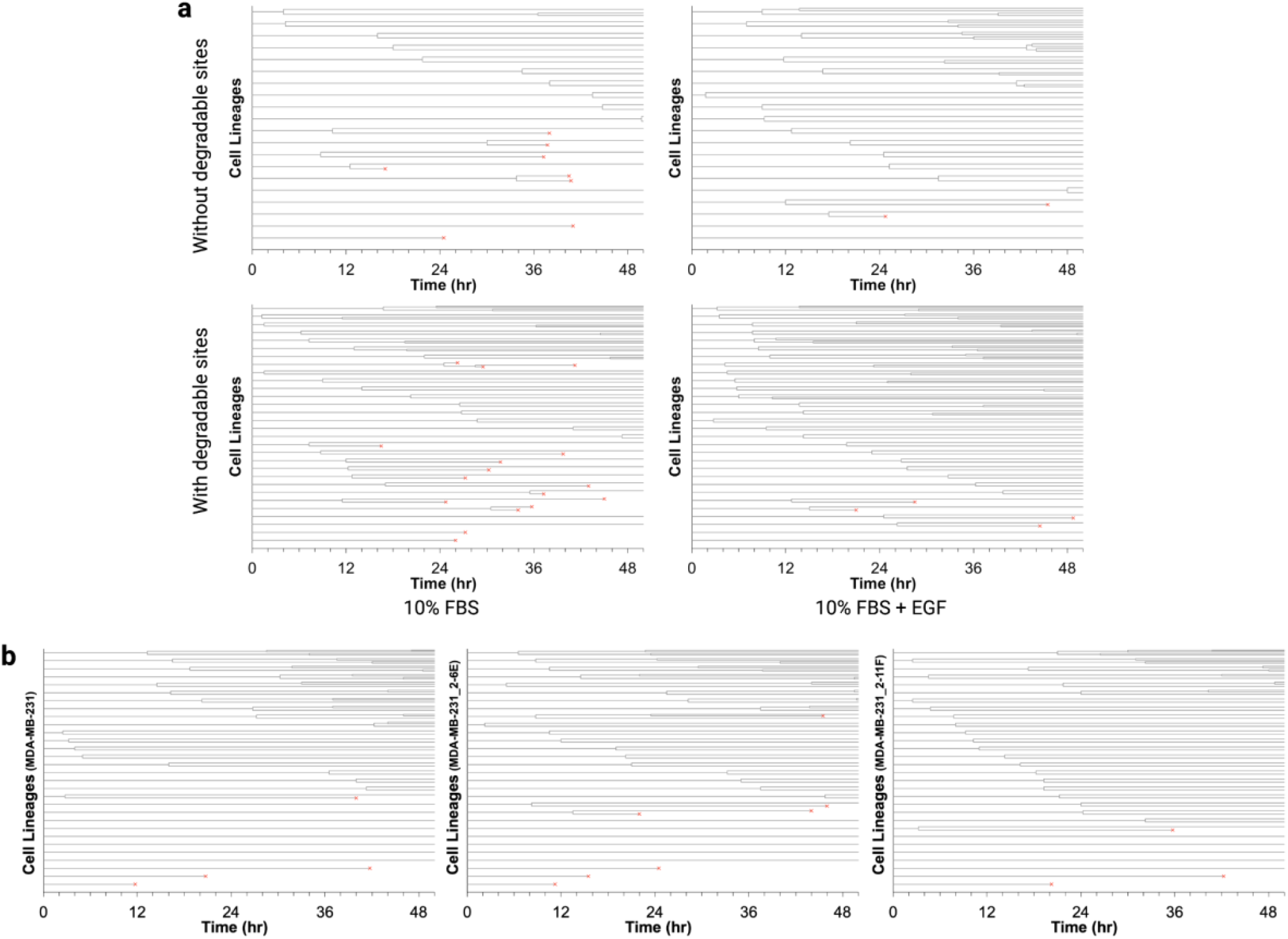
Cell lineage analysis for two TNBC cell lines, HCC1143 and MDA-MB-231, in a 3D gel environment. (**a**). Shows lineage analysis for triple negative breast cancer HCC1143 cells cultured in 3D PEG-RGD gels with or without MMP-degradable sites for 50 hours in 10% FBS containing condition (left column) and 10% FBS with EGF addition at 100ng/mL (right column). From the serum free condition, 20 cells were randomly selected for the cell lineage trees, and from the serum containing condition, 30 cells were randomly selected for the cell lineage trees. (**b**). Shows lineage analysis for triple negative breast cancer MDA-MB-231 parental cell line and its subclones. 30 cells were randomly selected for cell lineage analysis.

## Supplemental video

**S5. Dividing HCC1954 in 3D Collagen gel**.

This movie file shows a single HCC1954 cell seeded in 3D collagen gel with FBS-containing media is dividing at 16 hour 30 minute time point.

